# Pharmacogenetics at scale: An analysis of the UK Biobank

**DOI:** 10.1101/2020.05.30.125583

**Authors:** Greg McInnes, Adam Lavertu, Katrin Sangkuhl, Teri E. Klein, Michelle Whirl-Carrillo, Russ B. Altman

## Abstract

Pharmacogenetics (PGx) studies the influence of genetic variation on drug response. Clinically actionable associations inform guidelines created by the Clinical Pharmacogenetics Implementation Consortium (CPIC), but the broad impact of genetic variation on entire populations is not well-understood. We analyzed PGx allele and phenotype frequencies for 487,409 participants in the U.K. Biobank, the largest PGx study to date. For fourteen CPIC pharmacogenes known to influence human drug response, we find that 99.5% of individuals may have an atypical response to at least one drug; on average they may have an atypical response to 12 drugs. Non-European populations carry a greater frequency of variants that are predicted to be functionally deleterious; many of these are not captured by current PGx allele definitions. Strategies for detecting and interpreting rare variation will be critical for enabling broad application of pharmacogenetics.

## Introduction

Drug-based interventions play a primary role in medical treatment; more than 72% of visits to clinics and hospitals in the United States result in drug therapy^1^. An individual’s genetic makeup can have a profound impact on how they respond to many drugs. Therefore, the field of pharmacogenetics (PGx), is vital to improving modern medicine and prescribing practices^2^.

The practical value of PGx testing has increased as the field has discovered and characterized high impact haplotypes. These haplotypes are catalogued and named by PharmVar (www.pharmvar.org) using a nomenclature system typically based on “star alleles”^3–5^. Generally, the relationship between drug response and pharmacogenes is investigated through targeted studies on small groups of human subjects. The findings of these studies are aggregated through curation efforts such as PharmGKB (www.pharmgkb.org)^6^. The Clinical Pharmacogenetics Implementation Consortium (CPIC; cpicpgx.org) and other organizations assign a clinical function to star alleles based on published experimental research and create peer-reviewed and evidence-based clinical practice guidelines^7,8^. The clinical utility of PGx testing was recently recognized by UnitedHealthcare’s decision to extend coverage to PGx testing in the case of antidepressants and antipsychotic medications^9^.

PGx testing efforts are not yet capable of robustly handling rare genetic variation. Rare variants can be high impact, but are unlikely to be identified by a genotyping array or included in an established haplotype definition^10^. Most PGx testing in the US is currently implemented using genotyping arrays As a result, test results may be based on incomplete or partial allele definitions or proxy variants to assign PGx haplotypes, which may not represent the actual haplotype (as would be revealed by full and error-free sequencing) in the subject^11,12^. Developing more robust methods for assigning function to PGx haplotypes is an active area of research^13^. Unfortunately, the extent to which existing haplotypes definitions capture all important genetic variation within pharmacogenes is not well characterized^11,12,14^.

We used nearly 500,000 genotypes and 50,000 exome sequence samples in the UK Biobank to analyze pharmacogenetic variation at a population scale. To this end, we developed PGxPOP, a PGx matching engine, that is based on PharmCAT^15^ and uses its associated PGx allele definitions, to characterize pharmacogenetic allele and phenotype frequencies at scale. PGxPOP extends the capabilities of PharmCAT by generating diplotypes from population scale datasets^15^. Additionally, PGxPOP is built as a research tool; unlike PharmCAT, it does not create output for clinical implementation including patient-level reports containing genotype-based drug dosing recommendations. This study represents the largest study of pharmacogenetic allele and phenotype frequencies to date and investigates both the power and limitations of current star allele definitions. Our findings demonstrate the great value of characterizing allele frequencies in large populations, but highlights the need for more PGx research on historically under-studied populations, and the importance of using sequencing platforms that are capable of capturing rare genetic variation.

## Methods

### Data

This analysis focused on subjects enrolled in the UK Biobank, a prospective study of more than 500,000 individuals in the United Kingdom for whom detailed personal information, clinical data, and genetic data have been collected^29^. We use two sources of genomic data from the UK Biobank, genotype data imputed from sites collected on the Axiom Biobank Array (dataset release version 2), and exome sequencing data from the SPB pipeline (2/12/2020 rerelease of corrected data)^29,30^. We implemented several quality control measures to ensure high quality genetic data; we removed individuals from the analysis that were outliers based on heterozygosity and missing rates in the genetic data. These values are reported by the UK Biobank in the file “ukb_sqc_v2.txt”. Additionally, using VCFtools^31^, we excluded any loci that had a Hardy-Weinberg equilibrium p-value less than 1×10^−15^ or was missing in more than 10% of individuals. The imputed data was aligned to hg19 and the exome data was aligned to GRCh38. Each dataset was processed using the corresponding reference genomes. All data was phased using Eagle v2.4.1 (Fig. 1)^32^.

**Figure 1.**
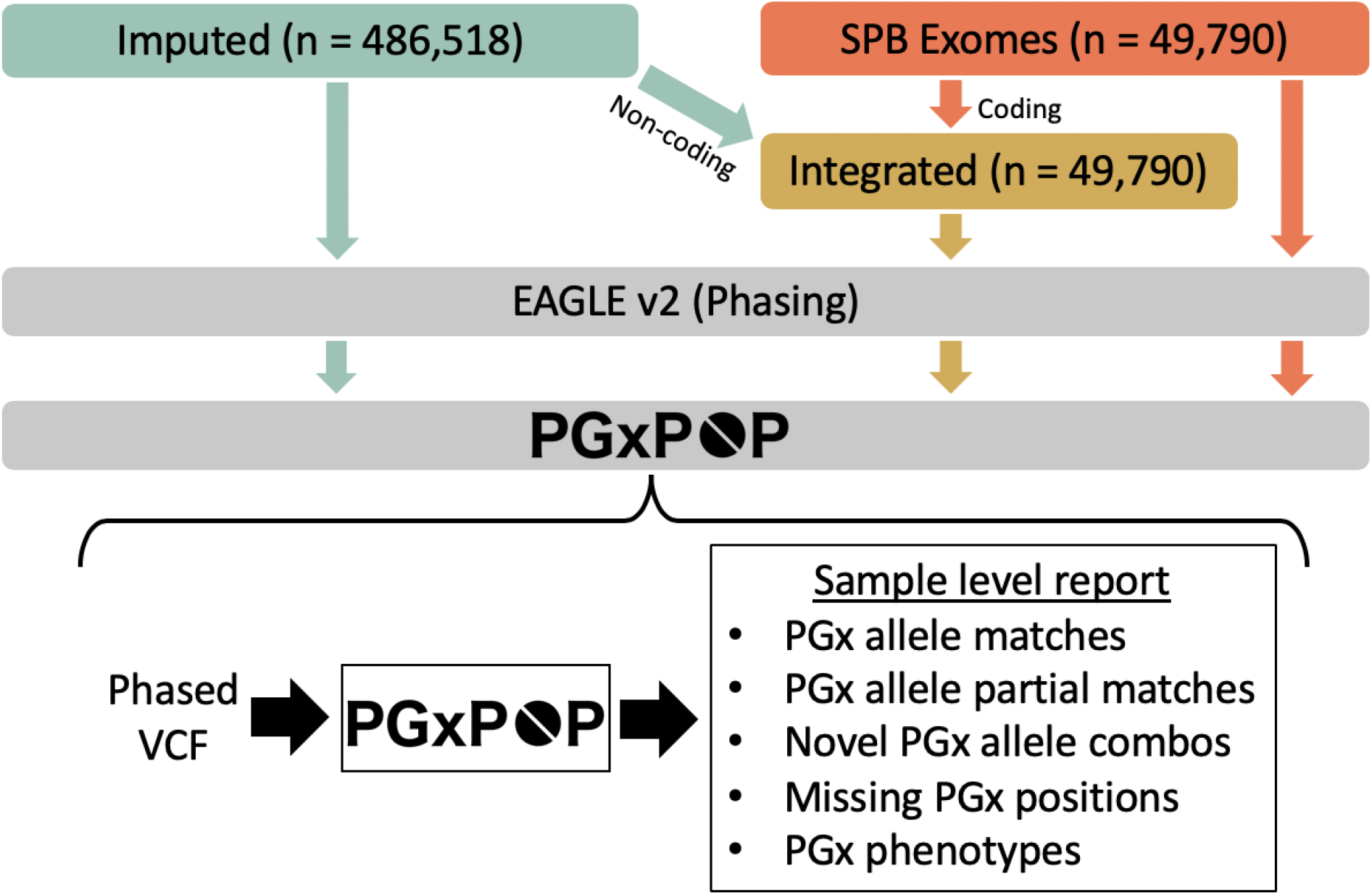
Analysis workflow. Our analysis comprises three data types, data imputed from genotype arrays, exome sequencing data, and an integrated call set that combines both. We phase all datasets using statistical phasing with EAGLE v2. We then generate pharmacogenetic alleles for all samples using PGxPOP and generate a report of the matching star allele, the variants contributing to that call, and the resulting phenotype.

We created an “integrated call set” by combining sequencing data of coding regions from the exome data and non-coding regions from the imputed genotype data. We used liftOver to map the variants in the imputed data to GRCh38^33^. Variants within the capture region on the exome array were extracted from the exomes and merged with intronic variants, and variants from 20 kilobases upstream and downstream of the transcription start and end sites (respectively). The newly merged variants were then phased with Eagle v2^32^.

We confirmed the ethnic population assignments using principal component analysis (PCA). We first group the self-reported ethnicity according to a standardized biogeographical system into African, European, East Asian, and South Asian^34^. Then, we calculated the mean and standard deviation of the first three principal components from a PCA of the genotype array data for each biogeographical group. Any sample whose reported ethnicity did not fall within three standard deviations of the mean for their reported ethnicity is referred to as “Other”.

### Cross-platform analysis

We analyzed the ability to call pharmacogenetic haplotypes and phenotypes for all fourteen genes (*CFTR, CYP2B6, CYP2C9, CYP2C19, CYP2D6, CYP3A5, CYP4F2, DPYD, IFNL3, NUDT15, SLCO1B1, TPMT, UGT1A1 and VKORC1*) across the three call sets: imputed, exome, and integrated. We limited the analysis to the 49,702 samples shared across all three platforms that pass quality control measures as described above. We calculated the diplotype and phenotype concordance between each platform and the integrated call set for each individual population and across all populations.

### Haplotype and phenotype calling

We developed PGxPOP, a tool that makes use of haplotype and phenotype definitions created by the PharmCAT effort^15^. We used the allele definition files in the PharmCAT GitHub repository (https://github.com/PharmGKB/PharmCAT), which are derived from allele definitions in PharmGKB^6^. PGxPOP reports exact matches to alleles as defined in PharmCAT’s allele definitions. In the case where one allele is completely subsumed within another (i.e. the defining variations for the allele are a proper subset of those for another allele), the allele with the maximum number of matching positions is reported. Importantly within this research setting, PGxPOP also reports partial matches or novel combinations of existing pharmacogenetic alleles (i.e. two distinct haplotypes on the same phased genotypes). In cases where there is a complete match to multiple haplotype definitions on the same strand, both haplotypes are reported in the program output with “+” notation when they are non-overlapping and “or” notation if there is overlap. For instance, if for *CYP2D6* both \**2* and \**9* alleles were found on the same strand, PGxPOP would report this as a \**2*+\**9* call, since the alleles for these two definitions are mutually exclusive. If instead, variants matching the \**35* and \**41* alleles were found on the same allele, where there is overlap at two positions, but there are also variants distinct to each, PGxPOP would report “*35+hg38:chr22.g.42127803C>T or *41+hg38:chr22.g.42130761C>T”, in order to represent all possible combinations of the alleles found at those positions. Found haplotypes are then mapped to predicted phenotypes based on published guidelines from PharmGKB and CPIC. PGxPOP was created as a research tool and is not intended for clinical use.

To enable analyses on large sample populations, we needed PGxPOP to process 100,000s of samples in several hours. We facilitated allele definition matching using matrix operations; PGxPOP computes the dot product of a reference allele matrix and the observed variant matrices (one for each phased haplotype in the VCF) and identifies matching haplotypes as those with a complete match to the haplotype definition across the largest number of positions (i.e. the sum of the dot product). In addition, we use tabix, a standard VCF indexing tool, to rapidly retrieve genomic data from compressed VCF files^35^.

We generated population specific haplotype, diplotype, and phenotype frequencies for the ethnic populations reported by UK Biobank. Haplotype and variant calls were generated across all samples for fourteen genes. We mapped sample diplotypes to phenotypes for all genes using CPIC guidelines, except *VKORC1* and *CYP4F2* because the CPIC guidelines do not provide that information for these genes. For phenotype prediction, the haplotypes were assigned the CPIC-associated function or activity values in cases of exact star allele matches. Phenotype was then determined based on the combination of the two alleles in the diplotype. For star allele combinations of alleles that included additional positions the allele function and phenotype was assigned as “not available”.

We use the star allele definitions when available for these genes with minor modifications. (1) We needed to make several assumptions about the ultimate phenotype of combination alleles and alleles carrying additional variants in order to assess the distribution of likely response phenotypes across the population. In these cases, we assume that if one of these alleles is nonfunctional, then the new combination of variants will not recover the function^36^. Thus, the alleles that include star alleles that result in no function are also assigned no function instead of ‘not available’. For example, if we identified a *CYP2D6* haplotype combination that includes *CYP2D6*4* and *CYP2D6*74* on the same strand (*CYP2D6*4+*74*), this haplotype would be determined to be “no function” even though function of *CYP2D6*74* is unknown. We do not extend this logic to alleles with decreased or increased function, except for *SLCO1B1* and *UGT1A1* where a haplotype carrying variants for a decreased function star allele is deemed to be decreased function. Additionally, any cystic fibrosis patient carrying a *CFTR* ivacaftor responsive allele is said to be ivacaftor responsive. (2) We modified the *SLCO1B1* allele definitions to exclude synonymous variants. We evaluated the ability to call star alleles in *SLCO1B1* with and without the three synonymous variants included in the existing star allele definitions. (3) For all INDELs we performed a search for identical INDELs in the sequencing data that may have been aligned differently. This was done by screening 50bp upstream and downstream of each INDEL in the definitions.

Importantly, structural variants were not called for *CYP2D6* or any other gene. Thus, we are not able to call star alleles with whole gene deletions (*CYP2D6*5*), duplications (e.g. *CYP2D6*1×2*), CYP2D7-2D6 hybrids (*CYP2D6*13*) or CYP2D6-2D7 hybrids including *CYP2D6*36*. This limits the assignment of CYP2D6 function and phenotypes since we are not able to determine CYP2D6 increased function alleles and therefore ultrarapid metabolizer and potentially miss no function alleles e.g. *CYP2D6*5, *13* or \**36*.

We calculated the burden of non-typical response phenotypes for each individual by counting the number of diplotypes with predicted non-typical response phenotypes across all twelve genes with phenotypes. Gene phenotypes were classified as “typical response” if they did not have any guidance away from a drug or its standard dosage based on all CPIC guidelines for that gene. Gene phenotypes were determined to have a non-typical response if any CPIC guidance recommended an alternate dosage or drug for that phenotype. Details of this heuristic can be found in Supplementary Table 1. We then determined the CPIC dosage recommendations for each subject for all 41 drugs with guidelines for any of the 14 genes in this study. This was done with PGxPOP using an encoding of the CPIC guidelines. For each drug, we determined the percent of the population that has been prescribed the drug by analyzing the general practice prescription data provided by the UK Biobank for more than 222,000 subjects. We considered any record of each drug (or a brand name version of the drug) being prescribed. We then calculated the percent of the population who had any record of being prescribed the drug.

### Deleterious variant analysis

In order to estimate the burden of deleterious variants in pharmacogenes we identified variants predicted to be deleterious in the exome data. We used a two-fold approach to predict if variants are deleterious. Any variant with a high IMPACT rating, such as frameshift indels, stop loss variants, was determined to be deleterious^37^. We then applied an ADME-optimized framework for predicting deleteriousness in pharmacogenes, which is an ensemble of deleteriousness prediction methods^38^. This approach enabled the prediction of the impact of missense variants as well as high impact variants. Variant IMPACT classes were determined using VEP^39^. All other annotations were generated using Annovar^40^.

Finally, we identified variants that were predicted to be deleterious that were not contained within existing star allele definitions. We calculated the aggregate deleterious variant allele frequency of all unaccounted-for deleterious variants by taking the sum of all allele counts for each deleterious variant not in an existing definition divided by the total number of samples.

## Results

### Platform concordance

We evaluated the concordance between three genomic call sets (imputed, exome, and integrated) for both diplotype and phenotype calling across twelve genes (Table 1). *IFNL3* and *VKORC1* were excluded from this analysis because the allele definition file for each gene consists of a single non-coding variant. For five genes where the majority of the variants of interest are in exons, we find very high (>96%) correlation between the integrated call set and both the imputed and exome call sets when calling both diplotypes and phenotypes (*CFTR, CYP2C9, TPMT, CYP4F2*, and *DPYD*). We observe a variety of concordance patterns for the other seven genes. For *CYP2C19*, which has a common non-coding variant upstream, the exome data is highly discordant with the integrated call set. Several genes have a mix of coding and non-coding variants, for these both platforms have low concordance with the integrated call set (*UGT1A1, CYP2D6, SLCO1B1*). For three genes, the exome data performs well, and the imputed data has lower concordance (*CYP2B6, CYP3A5*, and *NUDT15*). The imputed data for *NUDT15* has extremely low concordance with the integrated data; a variant that is rare in the population (rs746071566) was imputed for nearly all samples. Alluvial diagrams showing the change in haplotypes and phenotypes between the imputed and integrated call sets can be seen in Supplementary Figure 2.

**Table 1.** Platform concordance with integrated data is variable. We calculated the diplotype and phenotype concordance between the integrated call set and both contributing call sets, exome and imputed. For each gene we show the percent concordance (the percent of diplotypes or phenotypes that exactly match). Haplotypes for *IFNL3* and *VKORC1* contain only single variants that are in the non-coding regions, so the concordance is not listed for the exome data. *SLCO1B1* star alleles are determined excluding synonymous variants.

| Gene | Diplotype concordance w/ Integrated |  | Phenotype concordance w/ Integrated |  |
| --- | --- | --- | --- | --- |
|  | Imputed | Exome | Imputed | Exome |
| *NUDT15* | 0.01% | 99.63% | 0.04% | 99.67% |
| *UGT1A1* | 9.32% | 29.26% | 77.14% | 48.92% |
| *CYP2D6* | 34.23% | 84.50% | 64.86% | 86.64% |
| *CYP2B6* | 43.23% | 99.83% | 95.16% | 99.89% |
| *SLCO1B1* | 68.41% | 89.73% | 76.64% | 92.64% |
| *CYP3A5* | 85.48% | 100.00% | 85.69% | 100.00% |
| *CFTR* | 96.43% | 99.95% | 96.47% | 99.95% |
| *TPMT* | 97.76% | 99.93% | 99.17% | 99.93% |
| *CYP2C19* | 97.85% | 61.77% | 99.44% | 68.64% |
| *CYP2C9* | 98.23% | 99.85% | 98.36% | 99.86% |
| *CYP4F2* | 99.44% | 99.91% | 99.49% | 99.91% |
| *DPYD* | 99.60% | 95.67% | 99.61% | 95.68% |
| *IFNL3* | 1.00 | - | 1.00 | - |
| *VKORC1* | 1.00 | - | 1.00 | - |

We assessed the population-aware diplotype concordance between the imputed data and the integrated data in order to evaluate the population-specific accuracy of imputation. We find several genes for which there is a substantial decrease in imputation accuracy for some populations (Fig. 2). This gap is most extreme in *CYP3A5*, where subjects of European descent have a diplotype concordance of 86.8%, and subjects of African descent have a diplotype concordance of 14.7%. In total, four genes have a decrease of 10% diplotype concordance or more from the best performing ethnicity to the worst: *CYP3A5, CYP2B6, CYP2D6*, and *UGT1A1*.

**Figure 2.**
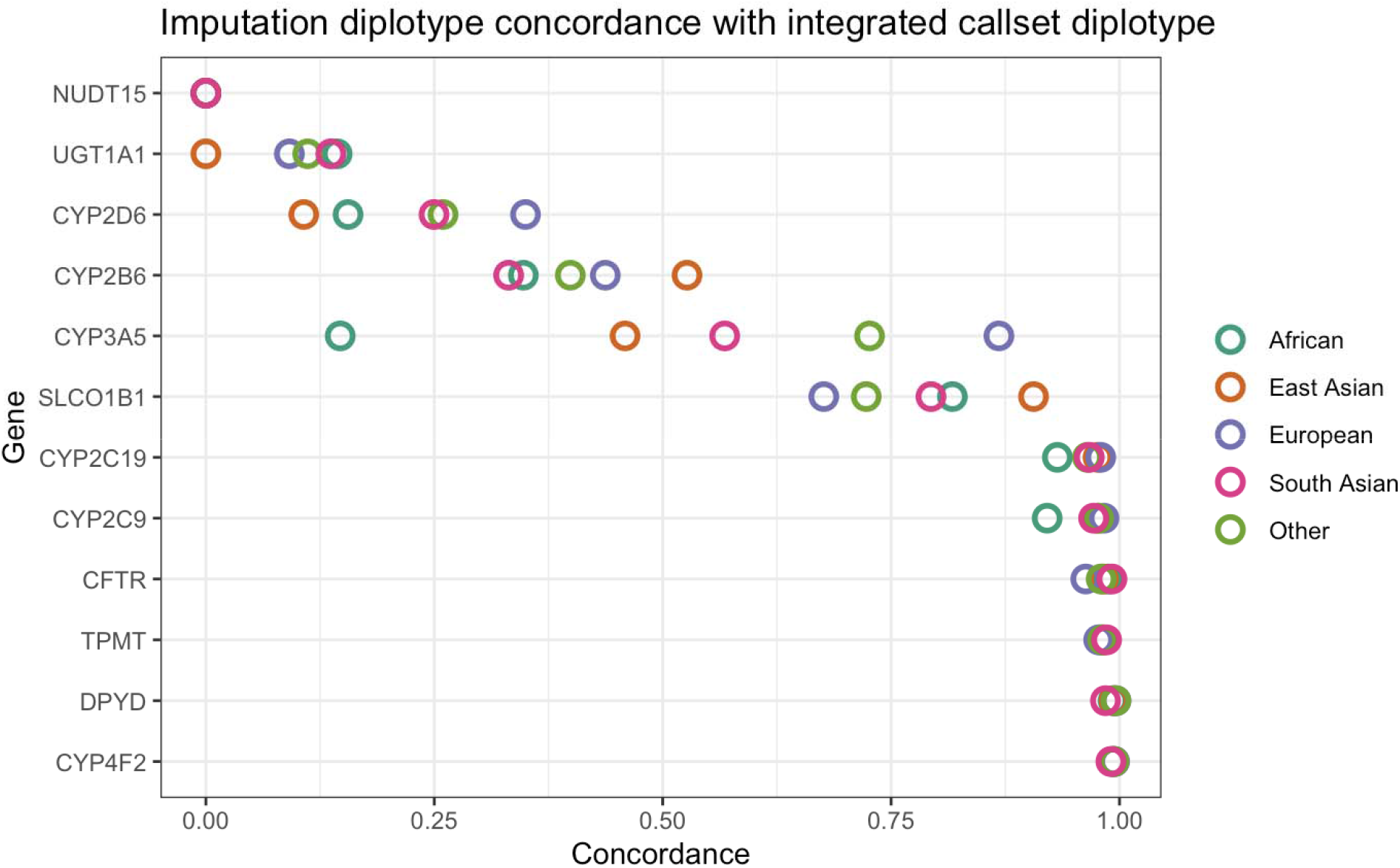
Concordance between diplotypes called from imputed data and integrated call set reveal inefficiencies in data imputed from genotype. The concordance is the proportion of diplotypes that exactly matched between the two call sets. We calculated population-specific concordance between the imputed data and integrated call sets. This comparison highlights the differences in the coding regions only, as the non-coding regions in the integrated call set are derived from the imputed data. Difference colors represent different global populations.

### Haplotype and phenotype calling

We analyzed haplotype and phenotype allele frequencies in clinically important pharmacogenes among individuals belonging to four global populations in the UK Biobank using a rapid haplotype matching engine. PGxPOP takes approximately six hours to call the diplotypes in all fourteen genes for the nearly 500,000 subjects. This analysis included 486,518 subjects with imputed data from genotyping arrays, 49,790 with exome sequencing data, and 49,790 subjects for whom an integrated call set was created by integrating the exome and non-coding regions from the imputed data. This study population includes subjects from four global populations (as well as 23,357 subjects who do not fall into a single population), verified using self-reported ethnicity and genetic ancestry (Table 2, Supplementary Figure 1). Haplotype and phenotype frequencies from the exome and integrated call sets for six cytochrome P450 genes included in our analysis are shown in Figure 3, and eight non-cytochrome genes in Figure 4. A full list of all haplotype, diplotype, and phenotype frequencies can be found for each call set in Supplementary file 1.

**Table 2.** Counts for each population analyzed and genetic data source. Subpopulations were grouped into global populations for broader analysis.

| Population | Subpopulation | Imputed count | Exome count | Integrated count |
| --- | --- | --- | --- | --- |
| European | British | 422,678 | 42,560 | 42,560 |
| European | Irish | 12,619 | 1,481 | 1,481 |
| European | Any other white background | 11,153 | 1,247 | 1,247 |
| European | White | 486 | 34 | 34 |
| European | Total | 446,936 | 45,322 | 45,322 |
| African | Caribbean | 4,110 | 632 | 632 |
| African | African | 3,089 | 329 | 329 |
| African | Any other Black background | 94 | 8 | 8 |
| African | Black or Black British | 22 | 3 | 3 |
| African | Total | 7,315 | 972 | 972 |
| South Asian | Indian | 5,533 | 682 | 682 |
| South Asian | Pakistani | 1,709 | 138 | 138 |
| South Asian | Bangladeshi | 212 | 16 | 16 |
| South Asian | Total | 7,454 | 836 | 836 |
| East Asian | Chinese | 1,456 | 170 | 170 |
| Other | Total | 23,357 | 2,490 | 2,490 |
| Total |  | 486,518 | 49,790 | 49,790 |

**Figure 3.**
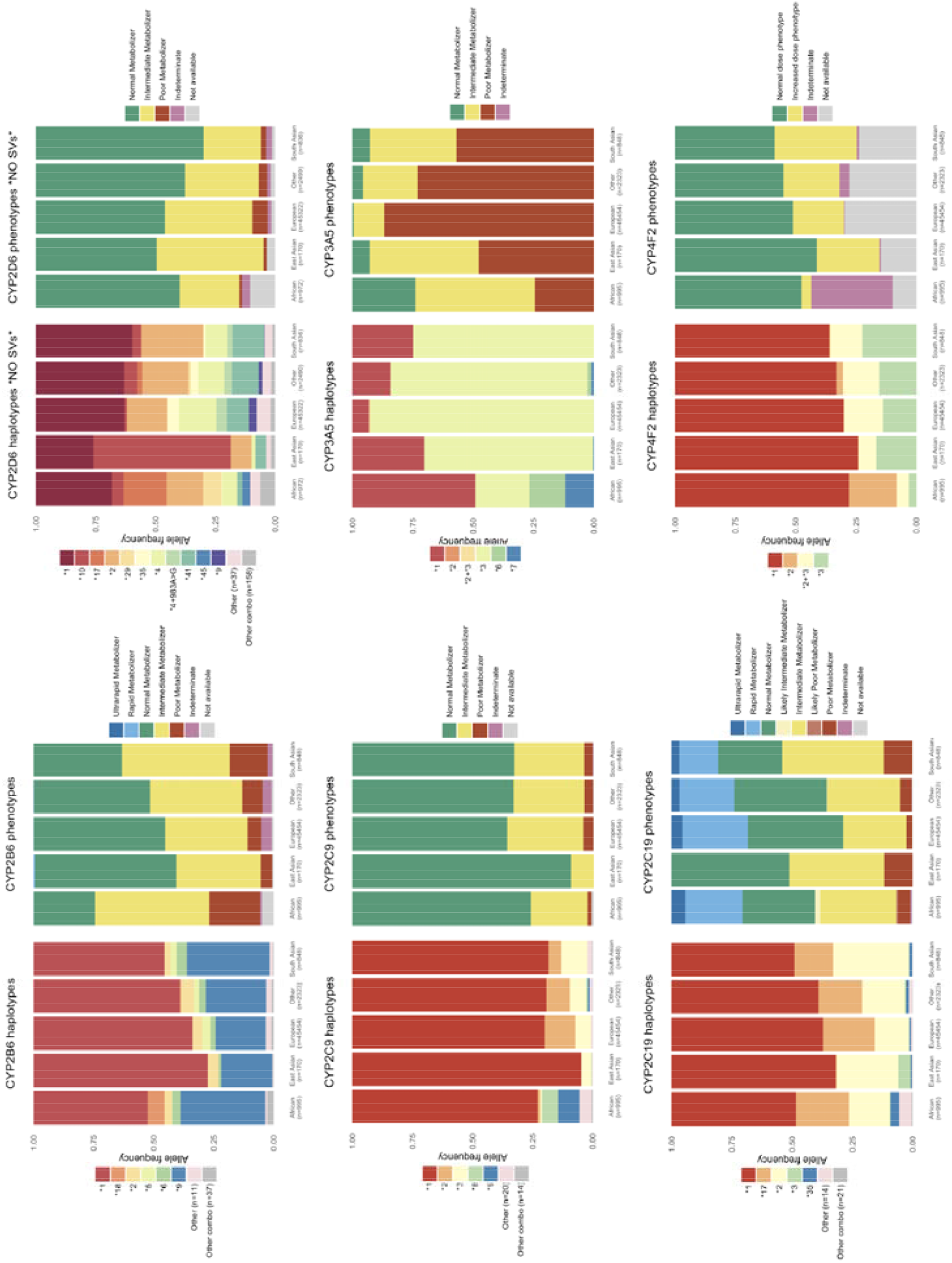
Star allele and phenotype frequencies for cytochrome P450 genes. Frequencies shown here are generated from the integrated call set which comprises nearly 50,000 subjects. The star allele frequency plots show all star alleles occurring with a frequency of 3% or greater. Any haplotypes with under 3% allele frequency in all populations are grouped into “Other”. Combination alleles, alleles that contain either partial or full matches of more than one star allele on the same strand occurring with less than 3% allele frequency are grouped in “Other combos”. The number of alleles in “Other” and “Other combos” is shown in the legend for each gene. Note that allele and phenotype frequencies for *CYP2D6* do not include structural variants.

**Figure 4.**
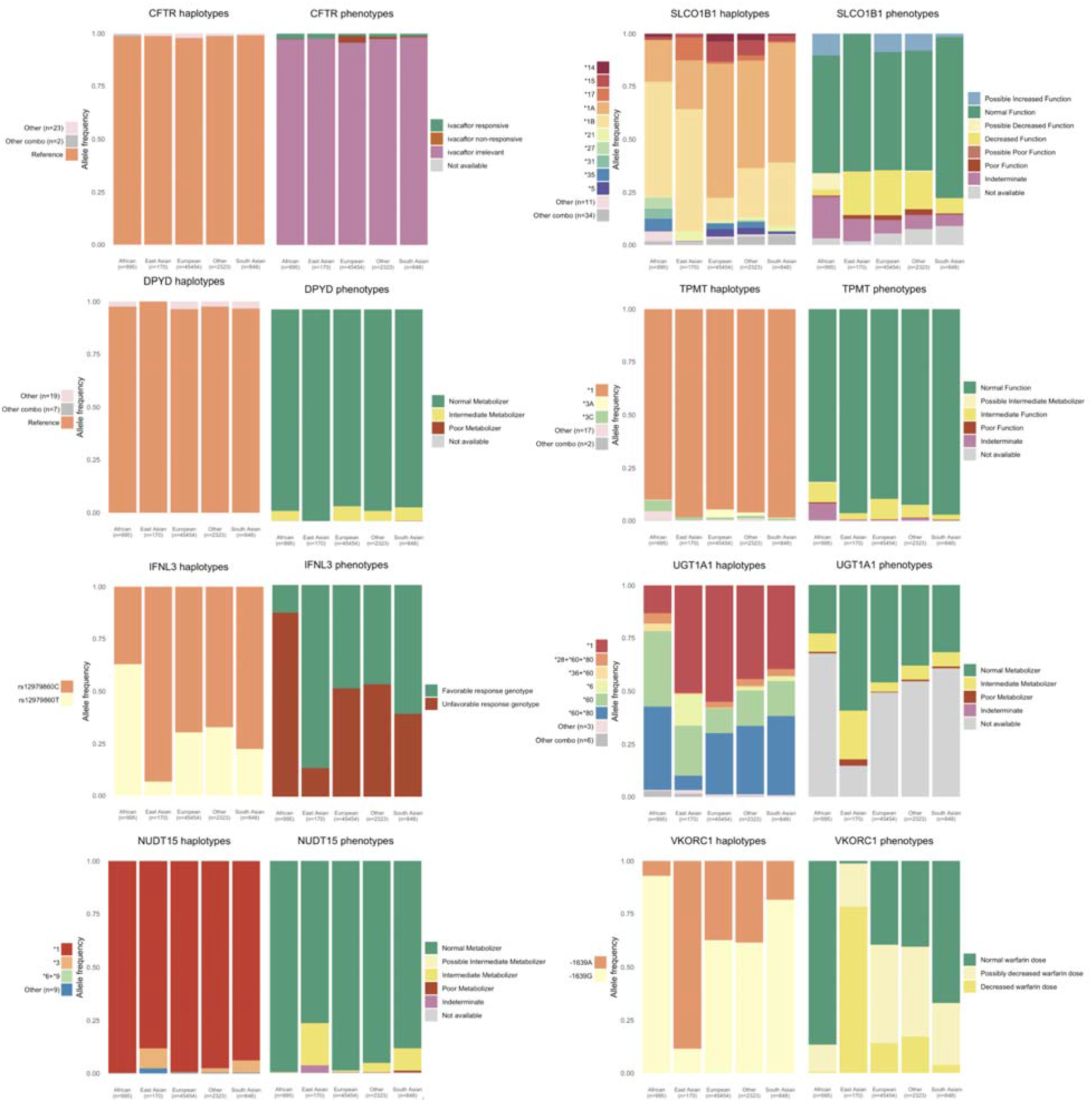
Star allele and phenotype frequencies for non-cytochrome P450 genes. Frequencies shown here are generated from the integrated call set which comprises nearly 50,000 subjects. The star allele frequency plots show all star alleles occurring with a frequency of 3% or greater. Any haplotypes with under 3% allele frequency in all populations are grouped into “Other”. Combination alleles, alleles that contain either partial or full matches of more than one star allele on the same strand occurring with less than 3% allele frequency are grouped in “Other combos”. The number of alleles in “Other” and “Other combos” is shown in the legend for each gene. *SLCO1B1* star alleles are determined excluding synonymous variants.

We find that on average subjects carry 3.7 non-typical response diplotypes for the fourteen pharmacogenes analyzed in the UK Biobank integrated call set, with 99.5% of subjects carrying at least one non-typical drug response diplotype (Fig. 5a). Subjects, on average, carry pharmacogene alleles that lead to atypical dosage guidance by CPIC for 12.2 drugs. We find for several frequently used drugs, a high number of people receive atypical dosage guidance, either recommended a different dose, different drug, or have a different recommended dosing procedure (Fig 5b). For example, simvastatin has been prescribed to 25% of the population, and 22.9 percent of all subjects carry either the rs4149056 C allele or *SLCO1B1* star alleles assigned possible decreased function (\**6*, \**9*, \**23*, \**31*), which indicates that a lower dose might be recommended due to increased risk of muscle toxicity^16^.

**Figure 5.**
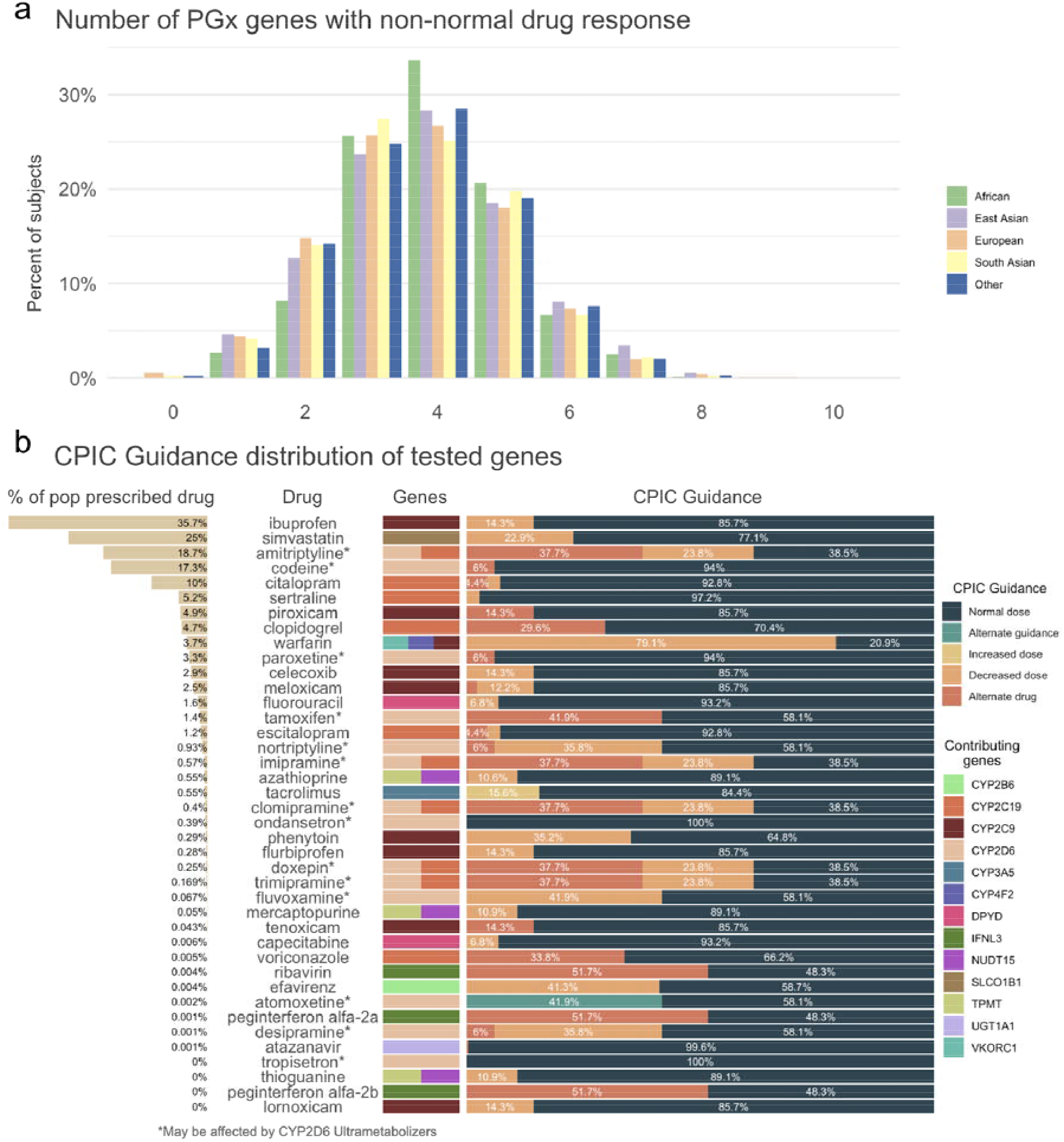
Frequency of pharmacogenes with a predicted non-typical response across the study population derived from the integrated call set and CPIC guideline recommendations for 41 drugs. a) The distribution of non-typical response alleles across each of the populations included in this study. Frequency of non-typical response pharmacogene alleles per subject range from 0 to 10, with a mean of 3.7. b) CPIC dosage guidance for 41 drugs that include recommendations based on any of the fourteen genes included in this study. We show the percent of the population that has ever been prescribed the drug, the drug name, the genes from this study that contribute to the recommendation, and the distribution of CPIC recommendations.

Star alleles with unknown or uncertain function, leading to an indeterminate phenotype, were found in nine genes. These are diplotypes where both haplotypes exactly match an existing star allele definition, but at least one of those haplotypes has unknown function. We find that 5.0% of subjects carry unknown or uncertain function star alleles in *SLCO1B1*, 4.2% in *CYP2B6*, and 1.7% in *CYP2D6*.

We find that for some genes, many novel combinations of alleles and allelic variants from existing allele definitions occur on a single haplotype in the integrated call set. These allele combinations can be a complete star allele or haplotype definition along with any number of additional variants from another previously defined allele. For example, 29.0% of the study population carries haplotypes that contain both the *CYP4F2*2* and *CYP4F2*3* variants on a single strand. Large numbers of novel allele combinations are also found in *CYP2D6* (159 unique combinations in 6.1% of subjects), *SLCO1B1* (34 in 2.9%), and *CYP2B6* (37 in 0.9%). At least one such allele combination was identified in twelve genes, the median number of allele combinations was eight, 288 were identified in total. *DPYD* and *CFTR* variation are represented by individual variants rather than star alleles, but combinations of variants were identified on a single strand for both genes. For analysis purposes, we assign function to these star allele and variant combinations by assuming that any no function star allele or variant will not be recovered by the addition of other variants in the same allele. The full details of the assumptions are described in the Methods. With these assumptions, predicted haplotype functions can be assigned to 102 of the variant combinations. The remaining 186 allele combinations cannot reliably be mapped to a function and are designated as ‘not available’ phenotype.

Individuals carrying a star allele with unknown or uncertain function or a star allele combination haplotype cannot be confidently mapped to a phenotype, and thus no CPIC drug dosing guidelines apply. Genes most impacted by this limitation are *CYP4F2* (30.2% of subjects), *SLCO1B1* (12.2%), *CYP2B6* (5.1%), and *CYP2D6* (3.4%). These counts exclude combination alleles for which we estimated function based on the rules defined in the previous paragraph.

We modified the *SLCO1B1* star allele definitions to exclude the three synonymous coding variants for the PGxPOP caller (chr12.g.21176827G>A, chr12.g.21178665T>C, and chr12.g.21178691C>T). These three variants appear in many combinations with the other core star allele variants and the star alleles that include these variants *18, *19, *20, *21 are assigned uncertain function. Including these three synonymous variants, 315 unique haplotypes were identified. The number of haplotypes decreased to 55 when those variants were removed. We find that when synonymous variants are included in the allele definition 77.9% of *SLCO1B1* haplotypes do not perfectly match one of the defined alleles and contain some combination of star allele variants and one or more variants from other definitions. This value drops to 2.9% when synonymous variants are excluded from the *SLCO1B1* definitions.

### Deleterious variant analysis

We estimated the burden of deleterious variants that are not currently included in allele definitions for eight of the fourteen genes in our study, *CYP2B6, CYP2C9, CYP2C19, CYP2D6, CYP3A5, NUDT15, SLCO1B1*, and *TPMT*. We predicted the deleteriousness of each variant found in the exome data and filtered out variants that were included in any existing allele definition, resulting in 478 deleterious variants across all eight genes (Fig. 6). Of the 478 deleterious variants identified, 244 have not been observed previously in gnomAD (Fig. 6c). All identified deleterious variants are rare (minor allele frequency < 1%). However, we find that 6.1% of all subjects carry at least one unaccounted for deleterious variant in one of these eight genes studied. To identify which populations are most underserved by current definitions we calculated the total frequency of all out-of-definitions deleterious variants in a population-specific manner (Fig. 6b). We find that across most genes, non-European populations carry the highest level of out-of-definition deleterious variants. For example, out-of-definition deleterious variants in *CYP2B6* have an allele frequency of 0.023 in the East Asian population. A full list of all identified deleterious variants can be found in Supplementary Table 3.

**Figure 6.**
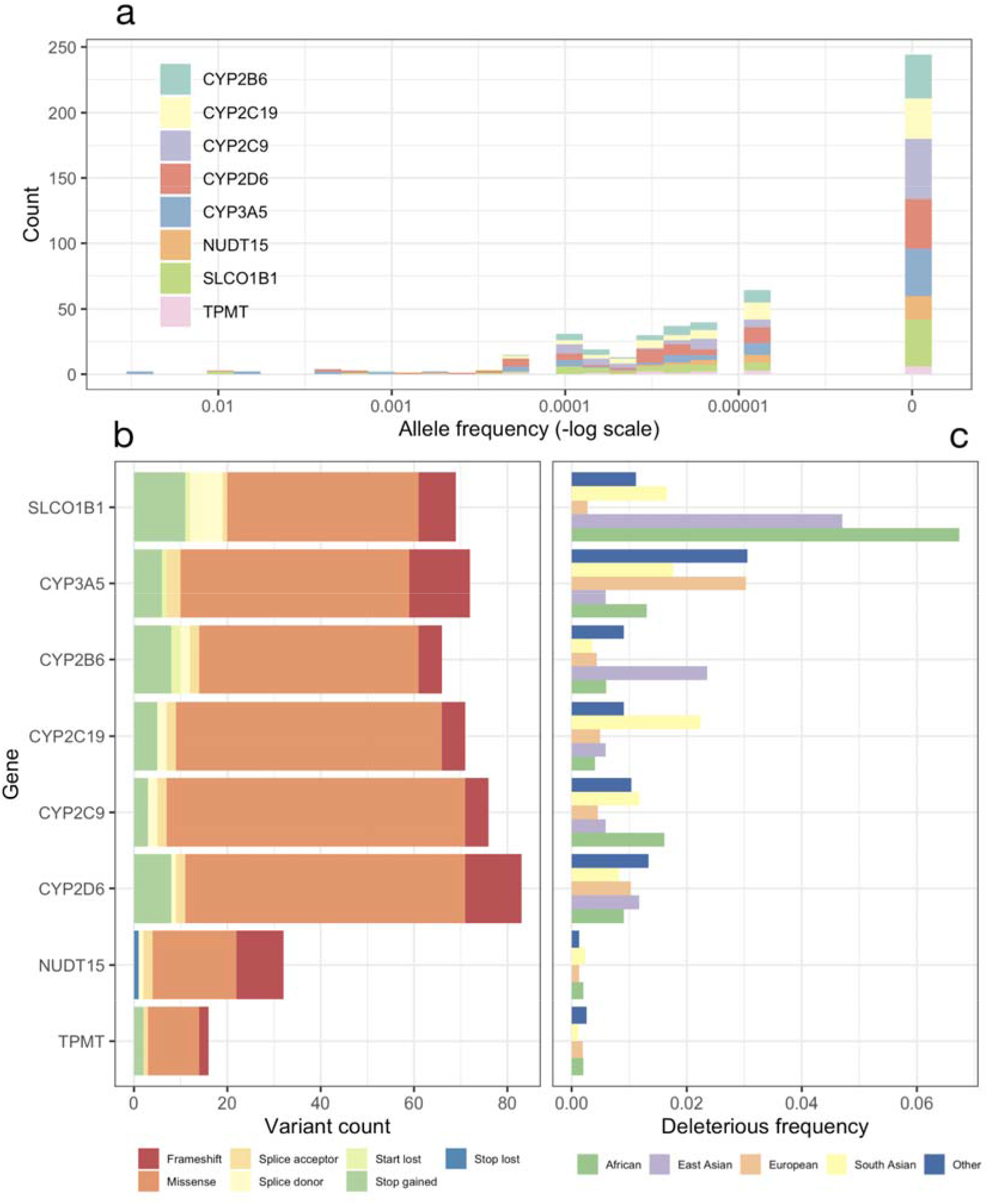
Analysis of deleterious variants not contained within existing star allele definitions. We identified presumptive deleterious variants in the exome sequencing data for eight genes by identifying probable loss of function variants as well as predicted deleterious missense variants. (a) shows the allele frequency of each probable deleterious variant in gnoMAD. Variants with an allele frequency of 0 were not identified in gnoMAD. (b) shows the number of deleterious variants identified as well as the frequency of each type of variant. (c) shows the total frequency of any deleterious variant in each population in the exome data. Concretely, the frequency represents the sum of allele frequencies for all deleterious variants not found within existing star allele definitions for each population.

## Discussion

Here we present a pharmacogenetic analysis of 487,409 participants in the UK Biobank, the largest study of its kind by an order of magnitude. The study cohort comprises mostly those of White British descent (n=442,615), however the minority populations in this study still represent the largest cohorts for those populations to date. Quantifying haplotype and phenotype frequencies at this scale enables a better understanding of the overall population risk of an adverse event when prescribing drugs related to these pharmacogenes, the coverage and accuracy of different genetic platforms, as well as the limitations of current pharmacogenetic allele definitions.

This analysis includes nearly 50,000 subjects with genetic data from both genotype array and exome data, providing an opportunity to assess the accuracy of each platform at a large scale. We find that for most genes there is high concordance between genomic data imputed from a genotyping array and sequencing data, for both haplotype and phenotype calls. Additionally, we show that the creation of an integrated call set, merging coding regions from exome data and non-coding regions from imputed data, leads to greater ability to identify haplotypes in genes that have functionally important non-coding variants (e.g. *CYP2C19*).

We find that several very important pharmacogenes are highly discordant between the imputed, exome, and integrated call sets, and some genes that have differences in imputation accuracy between populations--suggesting the need for great care when choosing a platform for pharmacogenetic analysis with regard to the gene and population of interest. For genes with splicing and other non-coding variants, exome data may not be sufficient (e.g. *CYP2C19*). While for highly polymorphic genes, imputed data may not be sufficient (e.g. *CYP2D6*). This highlights the potential clinical importance of having data from genome sequencing or a targeted capture array that includes coding and non-coding regions, such as PGRNseq^17^. Having genome sequencing would allow for the analysis of another crucial factor not captured by this study, the role of structural variants. For *CYP2D6* analysis, copy number variants and other structural aberrations are common and must be considered to make an accurate assessment of phenotype. Lack of structural variant analysis is a major limitation of this study’s ability to determine population level phenotype predictions of *CYP2D6*. However, we believe establishing star allele frequencies for star alleles identified from the variant data may still be useful.

Across all genes with haplotypes described by a star allele nomenclature, we find that there are haplotypes which are combinations of star allele variants that are currently not found together in any existing star allele definition. We also found combinations of individual variants in *DPYD* and *CFTR* on the same chromosome. Using array data can lead to the detection of only one of these alleles or variants, or the assumption that the alleles/variants are on different chromosomes. Either case can lead to the incorrect diplotype and phenotype assignment which could in turn result in an incorrect prescribing recommendation. We provide the star allele and variant combinations found in the UK Biobank population in the supplemental material to highlight the possibility and the frequency with which these occur.

The problem of accounting for rare variants is particularly challenging because numerous deleterious variants have not yet been submitted to PharmVar^3,4^, a resource devoted to cataloguing, defining and naming pharmacogene allele variation, so these deleterious variants do not contribute to current allele definitions. Individually, these variants are very rare; none have a minor allele frequency greater than 1%, and many of them are observed for the first time in this dataset. However, taken as a whole 17% of the population carries at least one deleterious variant in one of the fourteen genes analyzed that is not captured by the existing allele definitions. Deleterious variants within pharmacogenes are likely to have a strong effect on an individual’s PGx phenotype, indicating that 17% of the population in this study could benefit from a PGx guideline if one were to exist for their rare variation^14^. Non-European populations are the greatest affected, likely due to the European bias with which genetic studies have been conducted^18,19^. A greater effort to study pharmacogenetics in broader populations is necessary to make pharmacogenetics more accessible to the global community.

To date, *SLCO1B1* has not been included in PharmVar. Instead, *SLCO1B1* alleles *1a-*36 have been defined in 5 publications^16,20–23^. We find that the 37 star alleles for *SLCO1B1* are not commonly found as the only allelic variation for that gene. Only 22.1% of the *SLCO1B1* alleles from the UK Biobank exactly match the star allele definitions from these publications. Three synonymous coding variants (chr12.g.21176827G>A, chr12.g.21178665T>C, and chr12.g.21178691C>T) were the most commonly found with other star allele variants and removing them from the star allele definitions increased the allele matches to 97.1%. Further studies of the *SLCO1B1* haplotypes to confirm these findings in other populations would help inform if the current star allele definitions should be altered to exclude these three variants.

Our observation of individuals carrying combinations of PGx haplotypes and the rare nature of deleterious variants indicates that the current allele-based system would benefit from additional population-scale studies of PGx variation. Novel variation could then be incorporated into existing or new PGx allele definitions. However, our analysis demonstrates the limitations of the current PGx allele definitions; it is important that the community identify causal variants (and their mechanisms) so that reliance on specific LD structures can be reduced—a particularly important consideration in admixed populations, which are a virtually infinite source of haplotype diversity. An alternative to defining individual haplotypes and/or driving mutations, is to take a top down approach, in which regions of the gene or genes themselves are deemed essential, and any deleterious variant within essential components can be assigned an inferred phenotype. Recent work on the development of data-driven PGx phenotyping methods indicates that given enough data, it might be possible to move away from variant level rule-based systems and towards data-driven machine-learning models capable of robustly handling unobserved genetic variation^13,24,25^. The challenges posed by rare variation is likely to be a consistent issue for the current PGx system and will likely grow over time as genotyping gives way to genome sequencing and more populations are studied in detail revealing rarer and/or private mutations harbored by individuals.

We find that many individuals whose genotype does not match with an existing PGx definition are from populations that are underrepresented in PGx studies. So, there is a need to perform broad sequencing of global populations in order to enhance known pharmacogenetic variants across underrepresented populations. Underrepresented populations historically have low engagement in genetic medicine, in part due to fear of discrimination or lack of trust, among other barriers^26^. The genomic medicine community needs to continue to work to overcome these barriers and encourage population diversity in studies, submitting discovered pharmacogenetic variants and impact to PharmVar to better represent global populations.

One major limitation of this study is that we do not consider the effects of structural variants. Copy number variation and structural variation are well known to be functionally important in *CYP2D6* and relatively frequent phenomena. We attempted to perform copy number analysis of the exome data, but the existing tools for calling CNVs from exome data were found to be poorly maintained. We attempted to use tools built specifically to call *CYP2D6* structural variants but were because they were either not able to use exome data as input^27^, or require the reads to be aligned to hg19, which was computationally intractable^28^. Other studies have called CNVs from genotyping array intensities in the UK Biobank, but the observed frequencies of CNVs from array data are significantly different from those observed in genome sequencing data, calling the accuracy of these methods into question. Once the UK Biobank releases genome sequencing data, an analysis of structural variation in *CYP2D6* and other pharmacogenes will be a valuable contribution.

## Availability

PGxPOP is freely available and can be downloaded from https://github.com/PharmGKB/PGxPOP. All data used in the study can be obtained by applying to the UK Biobank for access.

## Acknowledgements

This research has been conducted using the UK Biobank Resource under Application Number 33722. We thank all the participants in the UK Biobank study. A.L. is supported by the National Science Foundation Graduate Research Fellowship (DGE – 1656518). G.M. is supported by the Big Data to Knowledge (BD2K) from the National Institutes of Health (T32 LM012409). K.S., M.W.C., T.E.K. and R.B.A are supported by NIH/National Institute of General Medical Sciences PharmGKB resource, (U24HG010615). R.B.A. is also supported by NIH GM102365. Most of the computing for this project was performed on the Sherlock cluster. We would like to thank Stanford University, the PharmGKB resource, and the Stanford Research Computing Center for providing the computational resources that contributed to these research results.

## Author Contributions

G.M. and A.L. conceived of the study, wrote PGxPOP, and performed the analysis. K.S., T.E.K. and M.W.C. provided guidance on PGxPOP and verified the output of the caller. R.B.A. oversaw the study. All authors helped with manuscript writing.

## References

1. 2016 NAMCS Summary Web Tables.

2. Lavertu, A. et al. Pharmacogenomics and big genomic data: from lab to clinic and back again. Hum. Mol. Genet. 27, R72–R78 (2018).

3. Gaedigk, A. et al. The Pharmacogene Variation (PharmVar) Consortium: Incorporation of the Human Cytochrome P450 (CYP) Allele Nomenclature Database. Clin. Pharmacol. Ther. 103, 399–401 (2018).

4. Gaedigk, A. et al. The Evolution of PharmVar. Clin. Pharmacol. Ther. 105, 29–32 (2019).

5. Robarge, J. D., Li, L., Desta, Z., Nguyen, A. & Flockhart, D. A. The star-allele nomenclature: retooling for translational genomics. Clin. Pharmacol. Ther. 82, 244–248 (2007).

6. Whirl-Carrillo, M. et al. Pharmacogenomics knowledge for personalized medicine. Clin. Pharmacol. Ther. 92, 414–417 (2012).

7. Relling, M. V. & Klein, T. E. CPIC: Clinical Pharmacogenetics Implementation Consortium of the Pharmacogenomics Research Network. Clin. Pharmacol. Ther. 89, 464–467 (2011).

8. Lunenburg, C. A. T. C. et al. Dutch Pharmacogenetics Working Group (DPWG) guideline for the gene-drug interaction of DPYD and fluoropyrimidines. Eur. J. Hum. Genet. 28, 508–517 (2020).

9. UnitedHealthcare Pharmacogenetic Testing. (2019).

10. Kozyra, M., Ingelman-Sundberg, M. & Lauschke, V. M. Rare genetic variants in cellular transporters, metabolic enzymes, and nuclear receptors can be important determinants of interindividual differences in drug response. Genet. Med. 19, 20–29 (2017).

11. Reisberg, S. et al. Translating genotype data of 44,000 biobank participants into clinical pharmacogenetic recommendations: challenges and solutions. Genet. Med. (2018) doi:10.1038/s41436-018-0337-5.

12. Caspar, S. M., Schneider, T., Meienberg, J. & Matyas, G. Added Value of Clinical Sequencing: WGS-Based Profiling of Pharmacogenes. Int. J. Mol. Sci. 21, (2020).

13. Lauschke, V. M. & Ingelman-Sundberg, M. Emerging strategies to bridge the gap between pharmacogenomic research and its clinical implementation. NPJ Genom Med 5, 9 (2020).

14. Ingelman-Sundberg, M., Mkrtchian, S., Zhou, Y. & Lauschke, V. M. Integrating rare genetic variants into pharmacogenetic drug response predictions. Hum. Genomics 12, 26 (2018).

15. Sangkuhl, K. et al. Pharmacogenomics Clinical Annotation Tool (PharmCAT). Clin. Pharmacol. Ther. 107, 203–210 (2020).

16. Ramsey, L. B. et al. Rare versus common variants in pharmacogenetics: SLCO1B1 variation and methotrexate disposition. Genome Res. 22, 1–8 (2012).

17. Gordon, A. S. et al. PGRNseq: a targeted capture sequencing panel for pharmacogenetic research and implementation. Pharmacogenet. Genomics 26, 161–168 (2016).

18. Martin, A. R. et al. Human Demographic History Impacts Genetic Risk Prediction across Diverse Populations. Am. J. Hum. Genet. 100, 635–649 (2017).

19. Sirugo, G., Williams, S. M. & Tishkoff, S. A. The Missing Diversity in Human Genetic Studies. Cell 177, 26–31 (2019).

20. Tirona, R. G., Leake, B. F., Merino, G. & Kim, R. B. Polymorphisms in OATP-C: identification of multiple allelic variants associated with altered transport activity among European- and African-Americans. J. Biol. Chem. 276, 35669–35675 (2001).

21. Nozawa, T. et al. Genetic polymorphisms of human organic anion transporters OATP-C (SLC21A6) and OATP-B (SLC21A9): allele frequencies in the Japanese population and functional analysis. J. Pharmacol. Exp. Ther. 302, 804–813 (2002).

22. Nishizato, Y. et al. Polymorphisms of OATP-C (SLC21A6) and OAT3 (SLC22A8) genes: consequences for pravastatin pharmacokinetics. Clin. Pharmacol. Ther. 73, 554–565 (2003).

23. Niemi, M. et al. High plasma pravastatin concentrations are associated with single nucleotide polymorphisms and haplotypes of organic anion transporting polypeptide-C (OATP-C, SLCO1B1). Pharmacogenetics 14, 429–440 (2004).

24. McInnes, G. et al. Transfer learning enables prediction of CYP2D6 haplotype function. bioRxiv 684357 (2020) doi:10.1101/684357.

25. van der Lee, M. et al. A unifying model to predict variable drug response for personalised medicine. bioRxiv 2020.03.02.967554 (2020) doi:10.1101/2020.03.02.967554.

26. Fisher, E. R. et al. The role of race and ethnicity in views toward and participation in genetic studies and precision medicine research in the United States: A systematic review of qualitative and quantitative studies. Mol Genet Genomic Med 8, e1099 (2020).

27. Twist, G. P. et al. Constellation: a tool for rapid, automated phenotype assignment of a highly polymorphic pharmacogene, CYP2D6, from whole-genome sequences. NPJ Genom Med 1, 15007 (2016).

28. Lee, S.-B. et al. Stargazer: a software tool for calling star alleles from next-generation sequencing data using CYP2D6 as a model. Genet. Med. 21, 361–372 (2019).

29. Sudlow, C. et al. UK biobank: an open access resource for identifying the causes of a wide range of complex diseases of middle and old age. PLoS Med. 12, e1001779 (2015).

30. Van Hout, C. V. et al. Whole exome sequencing and characterization of coding variation in 49,960 individuals in the UK Biobank. bioRxiv 572347 (2019) doi:10.1101/572347.

31. Danecek, P. et al. The variant call format and VCFtools. Bioinformatics 27, 2156–2158 (2011).

32. Loh, P.-R., Palamara, P. F. & Price, A. L. Fast and accurate long-range phasing in a UK Biobank cohort. Nat. Genet. 48, 811–816 (2016).

33. Haeussler, M. et al. The UCSC Genome Browser database: 2019 update. Nucleic Acids Res. 47, D853–D858 (2019).

34. Huddart, R. et al. Standardized Biogeographic Grouping System for Annotating Populations in Pharmacogenetic Research. Clin. Pharmacol. Ther. 105, 1256–1262 (2019).

35. Li, H. Tabix: fast retrieval of sequence features from generic TAB-delimited files. Bioinformatics 27, 718–719 (2011).

36. PharmVar. https://www.pharmvar.org/criteria.

37. Calculated consequences. https://m.ensembl.org/info/genome/variation/prediction/predicted_data.html.

38. Zhou, Y., Mkrtchian, S., Kumondai, M., Hiratsuka, M. & Lauschke, V. M. An optimized prediction framework to assess the functional impact of pharmacogenetic variants. Pharmacogenomics J. 19, 115–126 (2019).

39. McLaren, W. et al. The Ensembl Variant Effect Predictor. Genome Biol. 17, 122 (2016).

40. Wang, K., Li, M. & Hakonarson, H. ANNOVAR: functional annotation of genetic variants from high-throughput sequencing data. Nucleic Acids Res. 38, e164 (2010).

