## Supplemental Figures for "Pharmacogenetics at scale: An analysis of the UK Biobank"

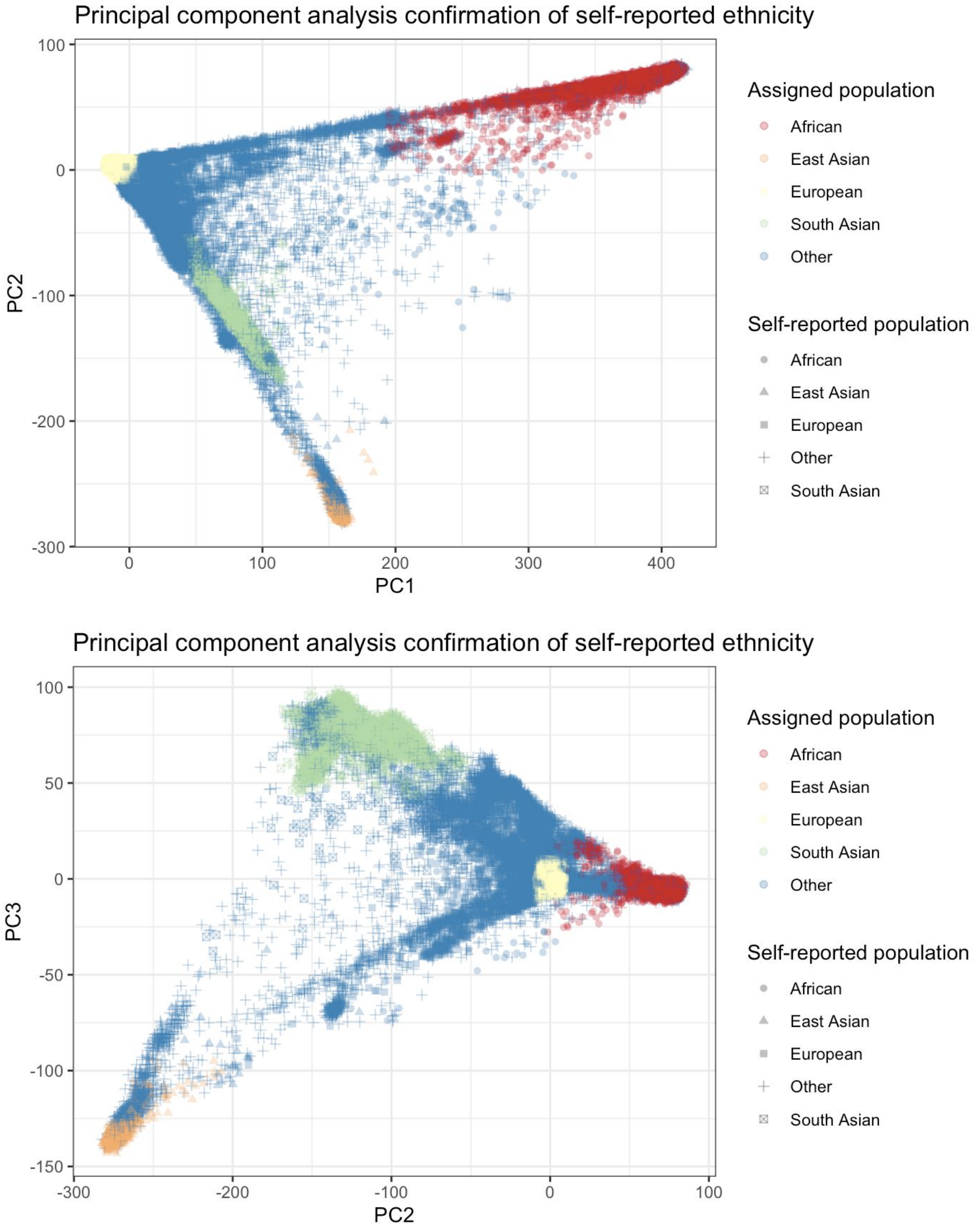


Supplementary Figure 1. Principal component confirmation of self-reported ethnicity. Ethnicity was confirmed using the first three principal components from PCA performed on genotype data. Genetic ancestry was assigned to each subject if they were within three standard deviations of the mean for the three principal components of their self-reported ethnicity. Any subject who's self-reported ethnicity did not match their genetic ancestry was marked as other.

Supplementary Table 2. Gene phenotypes leading to non-typical drug response as determined by CPIC

| Gene | Non-typical response phenotypes | Reference |
|---|---|---|
| CFTR | Ivacaftor non-responsive | 1 |
| CYP2B6 | Intermediate Metabolizer, Poor Metabolizer | 2 |
| CYP2C19 | Intermediate Metabolizer, Poor Metabolizer, Rapid Metabolizer, Ultrarapid Metabolizer | 3–6 |
| CYP2C9 | Intermediate Metabolizer, Poor Metabolizer | 7–9 |
| CYP2D6 | Intermediate Metabolizer, Poor Metabolizer, Ultrarapid Metabolizer* | 3,4,10–13 |
| CYP3A5 | Normal Metabolizer, Intermediate Metabolizer | 14 |
| CYP4F2 | Increased dose phenotype | 7 |
| DPYD | Intermediate Metabolizer, Poor Metabolizer | 15 |
| IFNL3 | Unfavorable response genotype | 16 |
| NUDT15 | Intermediate Metabolizer, Poor Metabolizer, Possible Intermediate Metabolizer | 17 |
| SLCO1B1 | Poor Function, Decreased Function, Possible Decreased Function | 18 |
| TPMT | Intermediate Function, Poor Function | 17 |
| UGT1A1 | Poor Metabolizer | 19 |
| VKORC1 | Decreased warfarin dose, possibly decreased warfarin dose | 7 |

### References


1. Clancy, J. P. *et al.* Clinical Pharmacogenetics Implementation Consortium (CPIC) guidelines for ivacaftor therapy in the context of CFTR genotype. *Clin. Pharmacol. Ther.* **95**, 592–597 (2014).
2. Desta, Z. *et al.* Clinical Pharmacogenetics Implementation Consortium (CPIC) Guideline for CYP2B6 and Efavirenz-Containing Antiretroviral Therapy. *Clin. Pharmacol. Ther.* **106**, 726–733 (2019).
3. Hicks, J. K. *et al.* Clinical pharmacogenetics implementation consortium guideline (CPIC) for CYP2D6 and CYP2C19 genotypes and dosing of tricyclic antidepressants: 2016 update. *Clin. Pharmacol. Ther.* **102**, 37–44 (2017).
4. Hicks, J. K. *et al.* Clinical Pharmacogenetics Implementation Consortium (CPIC) Guideline for CYP2D6 and CYP2C19 Genotypes and Dosing of Selective Serotonin Reuptake Inhibitors. *Clin. Pharmacol. Ther.* **98**, 127–134 (2015).
5. Scott, S. A. *et al.* Clinical Pharmacogenetics Implementation Consortium guidelines for CYP2C19 genotype and clopidogrel therapy: 2013 update. *Clin. Pharmacol. Ther.* **94**, 317–323 (2013).
6. Moriyama, B. *et al.* Clinical Pharmacogenetics Implementation Consortium (CPIC) Guidelines for CYP2C19 and Voriconazole Therapy. *Clin. Pharmacol. Ther.* **102**, 45–51 (2017).
7. Johnson, J. A. *et al.* Clinical Pharmacogenetics Implementation Consortium (CPIC) Guideline for Pharmacogenetics-Guided Warfarin Dosing: 2017 Update. *Clin. Pharmacol.*

*Ther.* **102**, 397–404 (2017).
8. Theken, K. N. *et al.* Clinical Pharmacogenetics Implementation Consortium Guideline (CPIC) for CYP2C9 and Nonsteroidal Anti-Inflammatory Drugs. *Clin. Pharmacol. Ther.* (2020) doi:10.1002/cpt.1830.
9. Caudle, K. E. *et al.* Clinical pharmacogenetics implementation consortium guidelines for CYP2C9 and HLA-B genotypes and phenytoin dosing. *Clin. Pharmacol. Ther.* **96**, 542–548 (2014).
10. Brown, J. T. *et al.* Clinical Pharmacogenetics Implementation Consortium Guideline for Cytochrome P450 (CYP)2D6 Genotype and Atomoxetine Therapy. *Clin. Pharmacol. Ther.* **106**, 94–102 (2019).
11. Crews, K. R. *et al.* Clinical Pharmacogenetics Implementation Consortium guidelines for cytochrome P450 2D6 genotype and codeine therapy: 2014 update. *Clin. Pharmacol. Ther.* **95**, 376–382 (2014).
12. Bell, G. C. *et al.* Clinical Pharmacogenetics Implementation Consortium (CPIC) guideline for CYP2D6 genotype and use of ondansetron and tropisetron. *Clin. Pharmacol. Ther.* **102**, 213–218 (2017).
13. Goetz, M. P. *et al.* Clinical Pharmacogenetics Implementation Consortium (CPIC) Guideline for CYP2D6 and Tamoxifen Therapy. *Clin. Pharmacol. Ther.* **103**, 770–777 (2018).
14. Birdwell, K. A. *et al.* Clinical Pharmacogenetics Implementation Consortium (CPIC) Guidelines for CYP3A5 Genotype and Tacrolimus Dosing. *Clin. Pharmacol. Ther.* **98**, 19–24 (2015).
15. Amstutz, U. *et al.* Clinical Pharmacogenetics Implementation Consortium (CPIC) Guideline for Dihydropyrimidine Dehydrogenase Genotype and Fluoropyrimidine Dosing: 2017 Update. *Clin. Pharmacol. Ther.* **103**, 210–216 (2018).
16. Muir, A. J. *et al.* Clinical Pharmacogenetics Implementation Consortium (CPIC) guidelines for IFNL3 (IL28B) genotype and PEG interferon-α-based regimens. *Clin. Pharmacol. Ther.* **95**, 141–146 (2014).
17. Relling, M. V. *et al.* Clinical Pharmacogenetics Implementation Consortium Guideline for Thiopurine Dosing Based on TPMT and NUDT15 Genotypes: 2018 Update. *Clin. Pharmacol. Ther.* **105**, 1095–1105 (2019).
18. Wilke, R. A. *et al.* The clinical pharmacogenomics implementation consortium: CPIC guideline for SLCO1B1 and simvastatin-induced myopathy. *Clin. Pharmacol. Ther.* **92**, 112–117 (2012).
19. Gammal, R. S. *et al.* Clinical Pharmacogenetics Implementation Consortium (CPIC) Guideline for UGT1A1 and Atazanavir Prescribing. *Clin. Pharmacol. Ther.* **99**, 363–369 (2016).

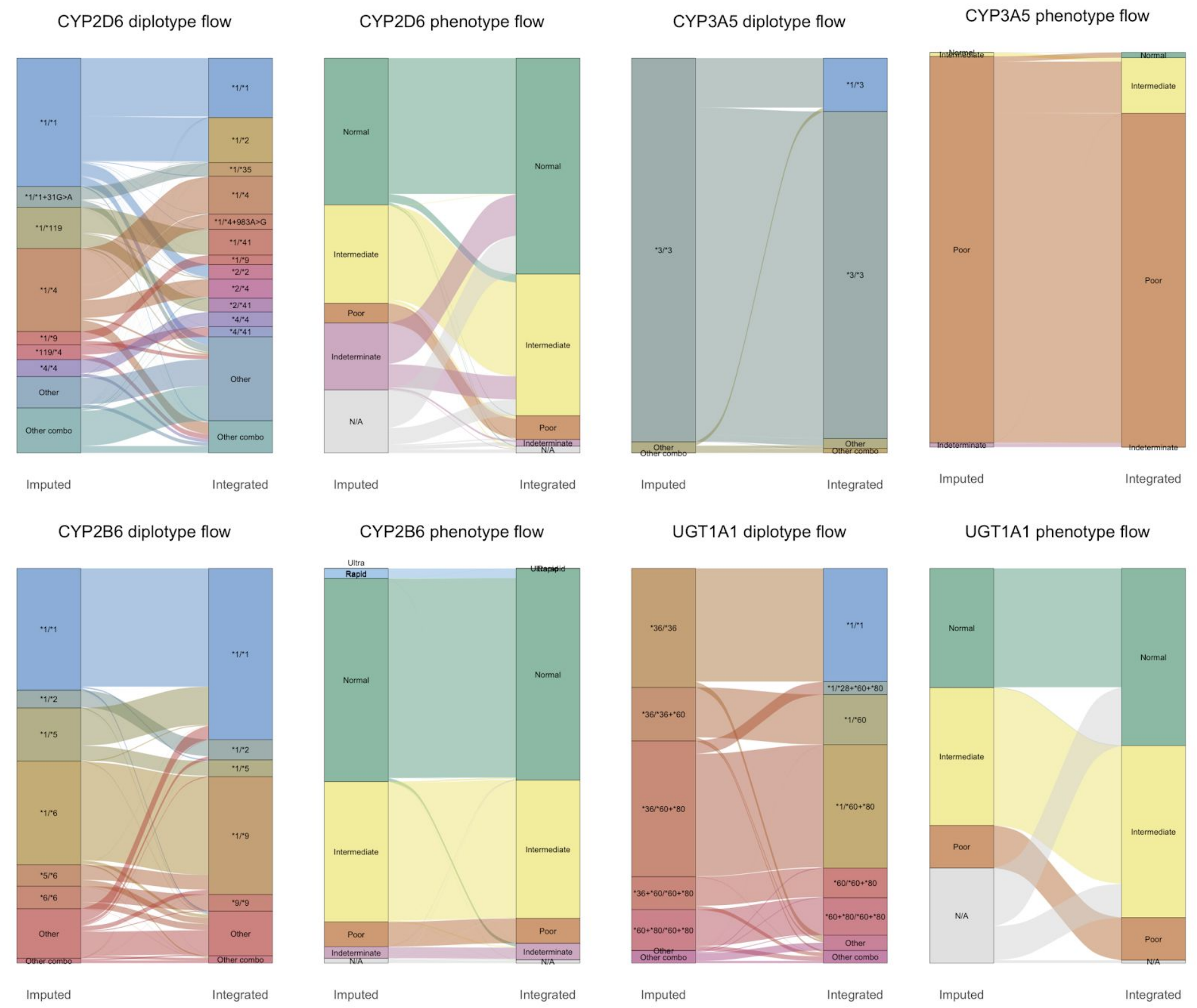


Supplementary Figure 2. Alluvial diagrams of diplotype and phenotype flow between platforms highlight frequent incorrect calls in the imputed data. We show diagrams of the four genes with the lowest concordance between the imputed and integrated callsets (excluding *NUDT15*). Diplotypes occurring with a frequency of less than 3% are grouped into “Other”, and diplotypes containing combination alleles occurring with less than 3% frequency are grouped into “Other combo”.
